# Recombinant production in *Escherichia coli* of functionally active α-hemolysin from the human pathogen *Staphylococcus aureus*

**DOI:** 10.1101/2025.01.14.632992

**Authors:** Jaime L. Díaz-Varela, Vittoria Sabia, Diego Heras-Márquez, Diego Laxalde-Fernández, Álvaro Martínez-del-Pozo, Sara García-Linares

**Affiliations:** Departamento de Bioquímica y Biología Molecular, Facultad de Ciencias Químicas, Universidad Complutense, 28040 Madrid, Spain

**Keywords:** Pore-forming proteins, nanodiscs, cholesterol, ITC, cytolysins, hemolysis

## Abstract

*Staphylococcus aureus* is a human opportunistic pathogen capable of causing multiple infections in both humans and animals. It secretes a group of exotoxins, known as hemolysins, which are released to enhance its pathogenicity. All of them exhibit cytolytic activity on a variety of host cell types, but α-hemolysin stands out for being the most thoroughly studied variant. In this work, we show the production and purification of *S. aureus* α-hemolysin following a straightforward protocol and in sufficient quantity to consider it as a potential procedure for future biotechnological approaches. Functional and structural characterization has indeed revealed that the protein is fully functional, confirming the key role of cholesterol in the necessary protein-lipid interaction. Furthermore, it has also been shown that the purified toxin can be assembled into single-particle individual pores within soluble lipid platforms in the form of cholesterol-containing nanodiscs.

## 1. Introduction

Cytotoxins are toxic agents capable of causing cellular damage, usually ending in cell death. Depending on their mechanism of action, they are categorized into diverse groups. For example, there are those that can alter the intracellular metabolism of their target cells and therefore need to cross the plasma membrane beforehand. Another group corresponds to toxins that exert their effect directly on that plasmatic membrane [1, 2]. Both groups of toxins have evolved to target this barrier, quite different among phyla, thus representing an opportunity for specificity. Proteins such as cholera or botulinum toxins belong to the first group. In the second one, toxic proteins like α-hemolysin (α-HL), produced by *Staphylococcus aureus*, stand out as prominent examples [1-3].

*S. aureus* is a human opportunistic pathogen capable of causing multiple infections in both humans and animals [4-6]. It is a Gram-positive bacterium that is part of the normal microbiota of 20-30 % of humans [6, 7]. Its mechanism of action is based on the synthesis of surface proteins, responsible for bacterial adherence to the host, and on the secretion of proteins that induce cell death and bacterial expansion [1]. The proteins secreted, such as coagulase or catalase, favor adhesion and survival of the microorganism. They function as aggregation factors, or by conferring protection against phagocytosis. A different group of secreted exotoxins, such as leukocidins or hemolysins, constitute another component of this arsenal, which in this case are released to enhance pathogenicity [8].

Within this latter group, four different hemolysins encoded by *S. aureus* have been described: α-, β-, γ-, and δ-HL. All of them exhibit hemolytic and cytolytic activity on a variety of host cell types [3], but α-HL is the most thoroughly studied variant. Accordingly, it is considered of foremost importance as a prototype of a β-pore-forming protein (PFP) [2, 9, 10]. PFPs are usually classified according to the nature of the secondary structure of the protein segments, or domains, which are inserted into the membrane. In the case of α-PFPs, they are constituted by α-helices [11, 12], whereas in β-PFPs pores are lined by β-sheets [2, 10]. α-HL is, in fact, a 33.2 kDa β-PFP consisting of 293 amino acids, with an important role in the virulence of the bacteria, due to its ability to form pores in the host cell membranes, leading to cell lysis [2]. In addition to this direct effect on the plasma membrane of its target cells, it has been also described that α-HL can exhibit immunomodulatory effects, potentially affecting the immune response of the infected organism [13].

As most PFPs [14], α-HL is synthesized as a water-soluble monomer that is able to form amphiphilic oligomers upon association with specific protein receptors in lipid bilayers. In this particular case, this metamorphic transformation results in a mushroom-shaped heptameric pore with a molecular weight of about 232 kDa, where three well-defined domains are distinguished: The cap, the stem, and the rim bort (Figure 1). *In vitro* studies have elucidated the mechanism of pore formation, starting from a water-soluble monomer, as well as the conformational changes observed during the process in the different domains [15].

**Figure 1.**
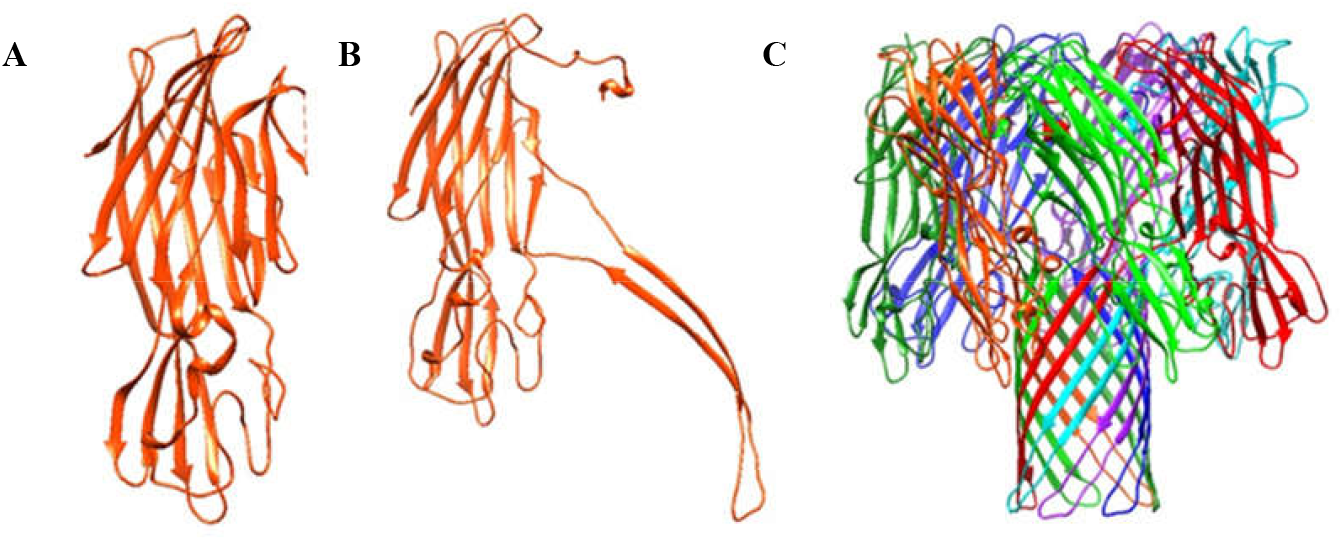
(A) α-HL structure of the water-soluble monomer (PDB 4YHD), (B) the pore protomer (PDB 7AHL), and (C) the heptameric pore structure (PDB 7AHL; each protomer is colored in one distinct color).

The pore structure is characterized by a transmembrane domain composed of 14 strands formed by seven β-hairpins, one from each protomer. These hairpins, rich in glycine residues, consist of alternating hydrophobic and hydrophilic residues, which, upon assembly, allow the hydrophobic residues to interact with the membrane and the hydrophilic ones to line the pore lumen [10, 14]. The pore has a diameter of 1.4 nm at its narrowest end and 4.6 nm for the most outer part of the structure (Figure 1C) [14].

Interaction with the lipid bilayer induces pore formation. Previous studies of the crystal structure of the protomer determined that dramatic conformational changes occur in the water-soluble structure of the monomers to form the final, thermodynamically stable, heptameric pore (Figure 1) [10]. In addition, these changes result in increased accessibility of the C-terminal domain, suggesting that it is projected towards the membrane surface and that it is important to achieve efficient oligomerization [16]. The detailed description of this molecular mechanism has been previously reported [15-19].

The interaction of α-HL with lipids, essential for its toxic activity, is a field that has not been characterized in depth yet. Studies made with multilamellar liposomes of different lipidic compositions have allowed the determination of the lipid components that enable and facilitate the interaction with α-HL. Phosphatidylcholine (PC) is required for α-HL binding and pore formation [20], with cholesterol (Chol) acting as an enhancer of the interaction, thus facilitating the interaction between the protein and the phosphocholine moiety of PC, as its presence leads to a more favorable orientation of this lipid headgroup [20, 21]. This role of Chol as an enhancer of the interaction has revealed that its presence in lipid membranes might also facilitate the interaction of α-HL with sphingomyelin (SM) [20], as has been described already for some other toxins [22]. This observation would fully agree with the well-known preference of Chol for SM [23]. However, even in the presence of Chol, the pore-forming activity of α-HL is still much more facilitated when interacting with PC than with SM [20].

Independently of its innate cytotoxicity against many eukaryotic cells, the ability of α-HL to generate pores in the plasma membrane has sparked interest because of its potential therapeutic and biotechnological applications [24, 25]. For example, its use as a therapeutic agent against some tumoral cells has been not only considered, but studied to some detail [26].

These studies relied on genetic engineering to modify pore specificity, and introducing the possibility of turning on and off the pore by means of the employment of a series of activators and inhibitors of different nature [27]. The goal was facilitating the penetration of proteases that would act on specific cancer cells. α-HL has also been considered as a potential facilitator for drug delivery. This study was based on transporting cargo within liposomes, and then inducing pore formation and subsequent release of the liposomal content caused by the interaction with a specific activator [27]. In recent years, the potential of PFPs as nanomachines has also been under investigation. This mechanism involves, for example, modification of protein residues to transform the pore into a nanoreactor [28, 29]. A good example of this approach is the creation of an α-HL nanopore that acts as an ATPase, a possibility which is already being studied [29]. In summary, the use of PFPs as a platform for new functions, whether for permeabilization or transportation of substances into various cell types, or for inducing enzymatic activities that facilitate the degradation of target molecules, holds significant interest [30]. The work presented herein shows the recombinant production and purification of α-HL, as well as the characterization of its lipid specificity and the assembly of the pore within water-soluble, Chol-containing, individual and thermodynamically stable platforms based on the employment of nanodiscs.

## 2. Materials and Methods

### 2.1. α-HL cloning in the plasmid pT7.7

The plasmid containing the DNA coding sequence of *S. aureus* α-HL was obtained from Integrated DNA Technologies (IDT). The primers detailed in Table 1 were designed for PCR amplification of α-HL cDNA including two restriction sites at the 5’ and 3’ termini for BamHI and NdeI. The amplification was performed in standard conditions using a Gene Amp PCR System 2400 Thermal Cycler (Perkin Elmer). The corresponding amplified fragment was purified using the PCR Preps DNA purification System kit (Promega), digested with the two enzymes mentioned and purified again using the same kit, after the corresponding separation using agarose electrophoresis. An identical procedure was used to obtain the plasmid pT7.7 fragment with the aimed BamHI and NdeI compatible ends. Ligation was made in standard conditions, using T4 DNA ligase (Biolab), followed by transformation of chemically competent *Escherichia coli* DH5αF’ cells allowed the selection of single bacteria colonies to propagate and purify the pT7.7α-HL final vector used for α-HL protein production. The correctness of the construction and the absence of unexpected mutation along the α-HL open reading frame was checked by Sanger DNA sequencing at the corresponding Complutense University facility.

**Table 1:**
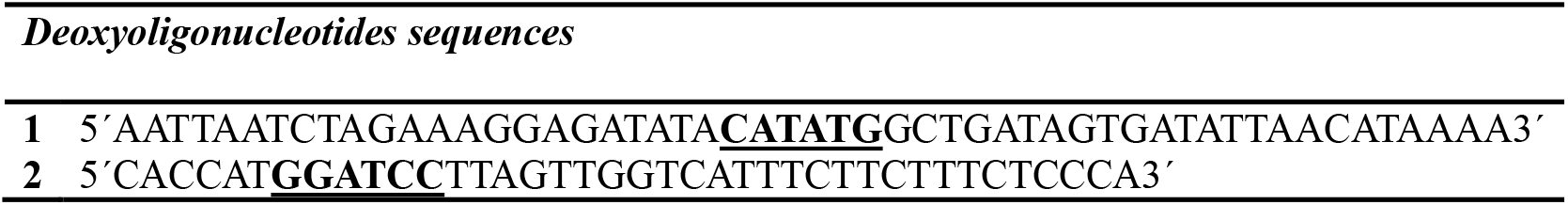
DNA sequence of deoxyoligonucleotides employed for obtaining the pT7.7α-HL plasmid. Restriction NdeI (1) and BamHI (2) sites appear in bold and underlined.

### 2.2. Protein production and purification

This pT7.7α-HL vector was then used for transformation chemically competent *E. coli* BL21(DE3)Gold using a Ca^2+^ and Rb^+^ based standard protocol, as described before [31, 32]. Small-scale production experiments were conducted to select the colony with the highest protein production efficiency. Once this was made, the selected colony was employed for larger scale production of the protein. In brief, this procedure was carried out in 2 L of LB containing Amp at 100 µg/mL at 37 °C and intense shaking (220 rpm) until an OD_600_ of 1.0 was reached.

Then, α-HL production was induced by adding isopropyl-β-D-thiogalactopyranoside (IPTG) at a final concentration of 1 mM and further incubation for 5 hours in identical conditions. After the induction period was finished, the culture medium was removed by centrifugation at 4700 g for 30 minutes at room temperature.

The cellular pellet was then resuspended in 50 mM sodium phosphate pH 7.0, containing 2 % (v/v) Tween 20, and subjected to sonication (7 cycles of 1 minute, 4 °C, 40 % amplitude). This cellular lysate was centrifuged for 30 minutes at 34700 g at 4 °C to discard the non-soluble cellular debris. The supernatant obtained was dialyzed (*cut-off* size 6-8 kDa) against 2.0 L of 10 mM sodium acetate, pH 5.0, 20 mM NaCl (“chromatography buffer”) to cause precipitation of *E. coli* self-proteins. These precipitated proteins were removed by centrifugation for 30 minutes at 34700 g at 4 °C. The dialyzed sample was then subjected to CM-52 ion exchange chromatography, equilibrated in the “chromatography buffer”. Elution was made with a salt linear gradient from 0.2 to 0.7 M NaCl. Fractions corresponding to α-HL were pooled, dialyzed against “chromatography buffer”, concentrated to 1.0 mg/mL, and stored frozen at -80 ºC. All protein samples were purified to homogeneity according to their electrophoretic behavior analyzed by 0.1 % (w/v) SDS−15 % (w/v) PAGE [33, 34].

### 2.3. Structural spectroscopic characterization

Far- and near-UV circular dichroism (CD) spectra were recorded using a Jasco 715 spectropolarimeter (Easton, MD). Fluorescence emission spectra were obtained on an SLM Aminco model 8000 spectrofluorimeter (Urbana, IL), equipped with a circulating water thermostat (Juliabo F30-C) to maintain a constant temperature. In both instances, the spectroscopic characterization was performed as previously described [35-38] in 10 mM sodium acetate, pH 5.0, 20 mM NaCl. Thermal denaturation was also evaluated as described [39]; registering the change in ellipticity at 220 nm as the temperature was raised at 0.5 ºC/min. T_m_ corresponds to the temperature at the midpoint of the transition observed.

Fluorescence emission spectra were recorded at 25 °C, between 280 and 400 nm, exciting at 275 and 295 nm, in 0.2 × 1.0 cm optical path quartz cells for the excitation and emission beams, respectively. The emission spectrum obtained upon excitation at 295 nm was normalized considering that, from 380 nm onwards, the contribution of Tyr is negligible, allowing the calculation of the Trp contribution. The contribution of Tyr was calculated as the difference between the emission spectrum excited at 275 nm and the normalized emission spectrum of Trp for that same wavelength. These measurements were also made as described [35-38], using 10 mM sodium acetate, pH 5.0, 20 mM NaCl.

### 2.4. Hemolysis assays

Hemolysis assays were performed in 96-multiwell plates as described [32, 37-40]. Briefly, erythrocytes from heparinized sheep blood (Dismalab) were washed in 10 mM Tris-HCl buffer, pH 7.4, 145 mM NaCl, to a final OD_655_ of 0.7 when mixing equal volumes of the cell suspension and buffer. Hemolysis was followed by recording the decrease in OD_655_ after addition of the erythrocyte suspension to different final concentrations of protein. A plate reader (Fluostar model 403) was employed to measure OD_655_. The value obtained with 0.1 % (w/v) Na_2_CO_3_ was considered as 100 % hemolysis. HC_50_ values were calculated as the concentration needed to get 50 % of hemolysis after 10 min of protein addition.

### 2.5. Lipid vesicles preparation

All lipids employed were obtained from Avanti Polar Lipids. Large unilamellar vesicles (LUVs) of DOPC:SM:Chol (1:1:1), DOPC:SM (4:1), DOPC:Chol (4:1) and DOPC were also prepared as described [36, 41-43]. Briefly, a phospholipid (0.1 – 1.0 mg) solution in 2:1 (v:v) chloroform:methanol was dried under a flow of nitrogen, and the dry film obtained was used to prepare a lipid dispersion by adding 0.5 - 2.0 mL of 10 mM Tris-HCl buffer, pH 7.4, 100 mM NaCl and 1 mM EDTA, briefly vortex mixing, and incubating for 1 h at 37 °C. This suspension of multilamellar vesicles was further subjected to fifteen cycles of extrusion at 37 °C through polycarbonate filters (100-nm pore size) (Nucleopore, Whatman) to obtain a homogeneous population of unilamellar vesicles. In order to quantify the lipid concentration, phosphorus titrations were performed following the adapted Bartlett method [44, 45].

### 2.6. Isothermal titration calorimetry

The interaction between α-HL and LUVs was quantitated by isothermal titration calorimetry (ITC), as previously described [37, 39, 46-49], using a VP-ITC calorimeter (Malvern MicroCal Worcestershire, U.K.). Solutions ranging between 4.0 - 5.0 μM protein concentrations were titrated by injection of 10 μL aliquots of lipid suspensions (phospholipid concentration of 2.5 mM) at a constant temperature of 37 °C. The buffer employed consisted of 10 mM Tris-HCl buffer, pH 7.4, 100 mM NaCl and 1 mM EDTA. Binding isotherms were adjusted to a model in which the protein binds to the membrane involving “N” lipid molecules [39].

### 2.7. SDS resistant oligomers

α-HL SDS-resistant oligomers were prepared following a protocol described before [50]. In brief, DOPC:SM:Chol (1:1:1) LUVs were incubated at various times (100-1100 seconds) and room temperature with the purified recombinant protein, at a lipid/protein molar ratio of 1000. The lipid vesicles were prepared in 10 mM sodium acetate, pH 5.0, 20 mM NaCl. The protein was dissolved in the same buffer but also containing 4% (w/v) SDS. The final concentration of SDS in the incubation mixture was 2.0% (w/v). At the indicated times, 10 µL aliquots were taken and analyzed on a standard 0.1% SDS-10% PAGE analysis.

### 2.8. Reconstitution of a soluble α-HL transmembrane pore on lipidic nanodiscs

Pore assembly was performed to mimic the actual situation encountered by α-HL in nature. Thus, instead of reconstituting the pore simultaneously with the assembly of nanodisc particles, the nanopores were prepared as described before for some other PFP [28]. Nanodiscs were first reconstituted [51, 52] from lipid stocks of DOPC:SM:Chol (1:1:1), which had been prepared at 50 mM lipid concentration in chloroform to prepare the convenient dried lipid aliquots. Chloroform was evaporated under a stream of nitrogen. The tubes were then placed in a desiccator for 3 hours under a high vacuum to remove all traces of the organic solvent. A pH 8.0 buffer containing 200 mM sodium cholate was then employed to rehydrate the lipid film. Typically, cholate was added up to twofold the lipid concentration. The tubes were then vigorously vortexed, heated at 60 °C for 10 min, and sonicated for 15 min. Once made, the MSP1E3D1 membrane scaffold protein was added to the lipid:detergent mixture to a final molar protein/lipid ratio of approximately 1/100 and incubated at room temperature for 30 minutes. Following extensive dialysis, the detergent was removed against 20 mM Tris-HCl buffer, pH 7.4, 100 mM NaCl and 0.5 mM EDTA, a chelating agent used only during the first dialysis treatment. The final dialyzed solution was centrifuged. The resulting clear supernatant was loaded onto a calibrated Superdex 200 column (Cytiva, Uppsala, Sweden), equilibrated in the same buffer employed for the dialysis, and attached to an ÄKTA purifier fast performance liquid chromatography (FPLC) system (Amersham Biosciences, Amersham, UK). The resulting chromatogram yielded a symmetrical peak corresponding to the expected nanodisc molecular size, which was then pooled and labelled as *purified empty nanodiscs*. α-HL was then mixed with this preparation at a α-HL/MSP1E3D1 molar ratio of 10/1. This mixture was incubated at room temperature for 1-2 hours and then used to perform electron microscopy analysis. The buffer employed here was 20 mM Tris-HCl buffer, pH 7.4, 100 mM NaCl.

Electron microscopy was performed on a JEOL Jem1400 transmission electron microscope operated at 100 kV at the National Centre for Electron Microscopy (CNME) located within the Complutense University campus. For sample visualization, 200-hole copper grids coated with a polymeric film were used and negative staining with 2% uranyl acetate (Panreac) was conducted.

## 3. Results and Discussion

### 3.1. α-HL purification

The soluble protein fraction remaining after the dialysis treatment (see the Methods section) was subjected to further fractionation by means of CM-52 ion exchange chromatography. The chromatogram obtained (Figure 2) revealed the presence of two well-resolved peaks. Spectroscopic and SDS-PAGE analysis of the fractions revealed that the second one corresponded to homogeneous isolated α-HL (Figure 3). The band observed in the gel, with an electrophoretic mobility corresponding to the expected molecular weight of 33.2 kDa, was estimated to be at least 95% homogenous.

**Figure 2.**
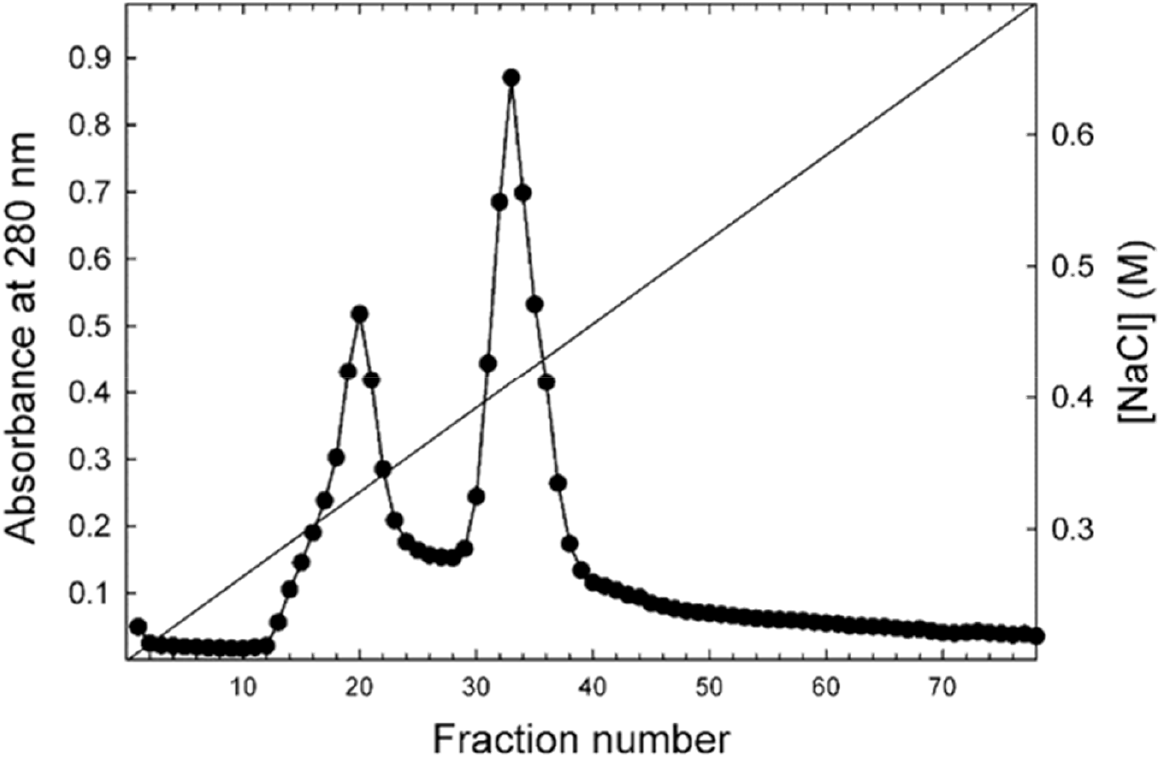
Chromatogram resulting from the CM-52 ion exchange chromatography purification step. The buffer employed was 10 mM sodium acetate, pH 5.0, 20 mM NaCl. Elution was made using a linear NaCl concentration gradient (0.2-0.7 M) which progression is also shown. The second peak corresponds to α-HL containing fractions.

**Figure 3.**
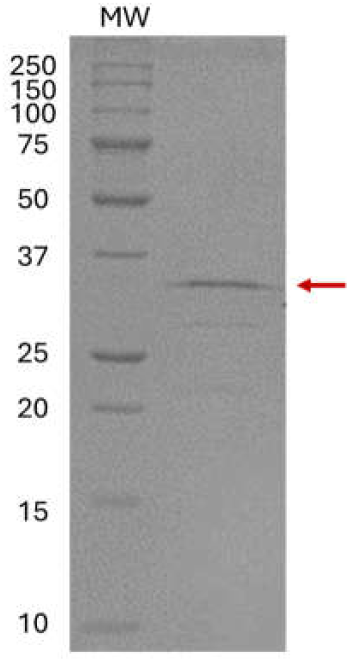
SDS-PAGE of a molecular weight (MW) protein standard (*left*) and the α-HL pool corresponding to the purified protein (*right*).

### 3.2. Spectroscopic characterization

The far-UV CD spectrum of purified α-HL (Figure 4A) was fully compatible with the β-sheet rich protein expected (Figure 1), with a minimum centered around 215 nm. The near-UV CD spectrum is also shown (Figure 4B). Both spectra together are compatible with a folded globular protein.

**Figure 4.**
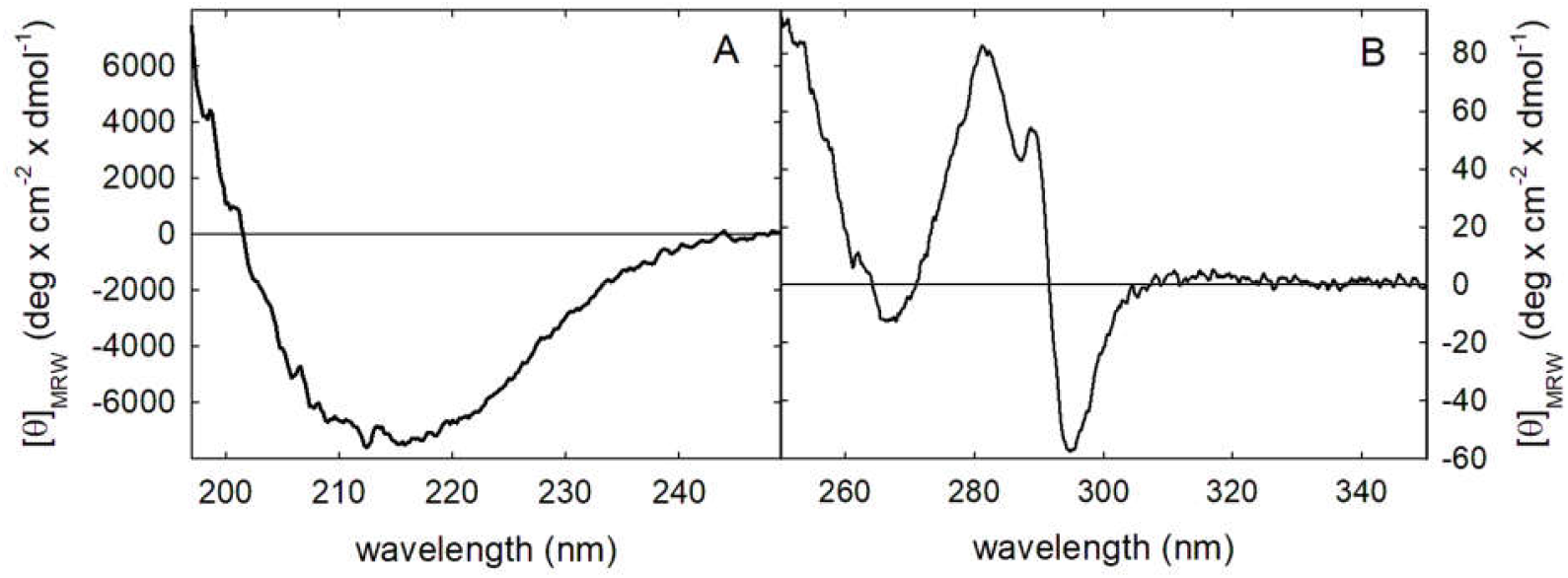
(A) Far-UV and (B) near-UV CD spectra of purified recombinant α-HL.

Regarding the thermal denaturation of the protein, Figure 5 shows a monophasic transition with a T_m_ value at pH 5.0 of around 53 ºC. Again, this all-none transition is also compatible with a stably folded globular protein. The T_m_ value is indeed in good agreement with the estimations made by other authors using differential scanning calorimetry [53].

**Figure 5.**
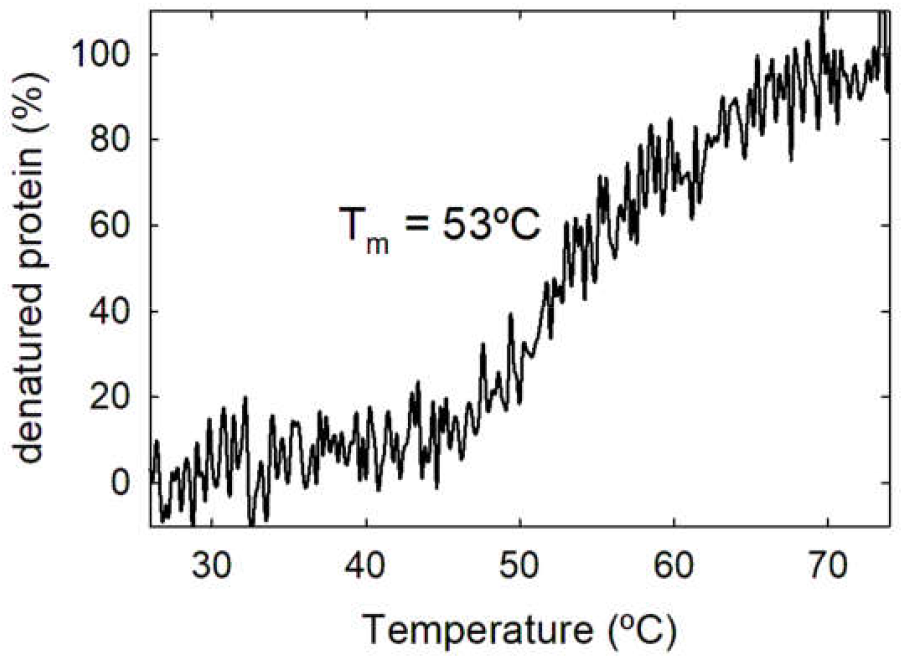
Purified recombinant α-HL thermal denaturation profile. The T_m_ value of 53 ºC corresponds to the temperature at the midpoint of the monophasic thermal transition. The buffer used was 10 mM sodium acetate, pH 5.0, 20 mM NaCl.

Fluorescence emission spectra for α-HL are shown in Figure 6. In addition to a shoulder around 300 nm, corresponding to its 14 Tyr residues, it displays a broad maximum centered at 325 nm. The width and heterogeneity of this maximum suggest quite different, but hydrophobic, microenvironments for the eight Trp residues in α-HL, as expected. This observation is corroborated by the calculated Tyr contribution spectrum, which does not correspond to the spectrum expected for only Tyr emission, revealing Trp emission heterogeneity that impedes its correct deconvolution. Overall, these fluorescence emission spectra also agree with the adoption of a compact globular structure, fully compatible with the CD spectra (Figure 4) and with the crystalline three-dimensional structure of monomeric water-soluble α-HL (Figure 1).

**Figure 6.**
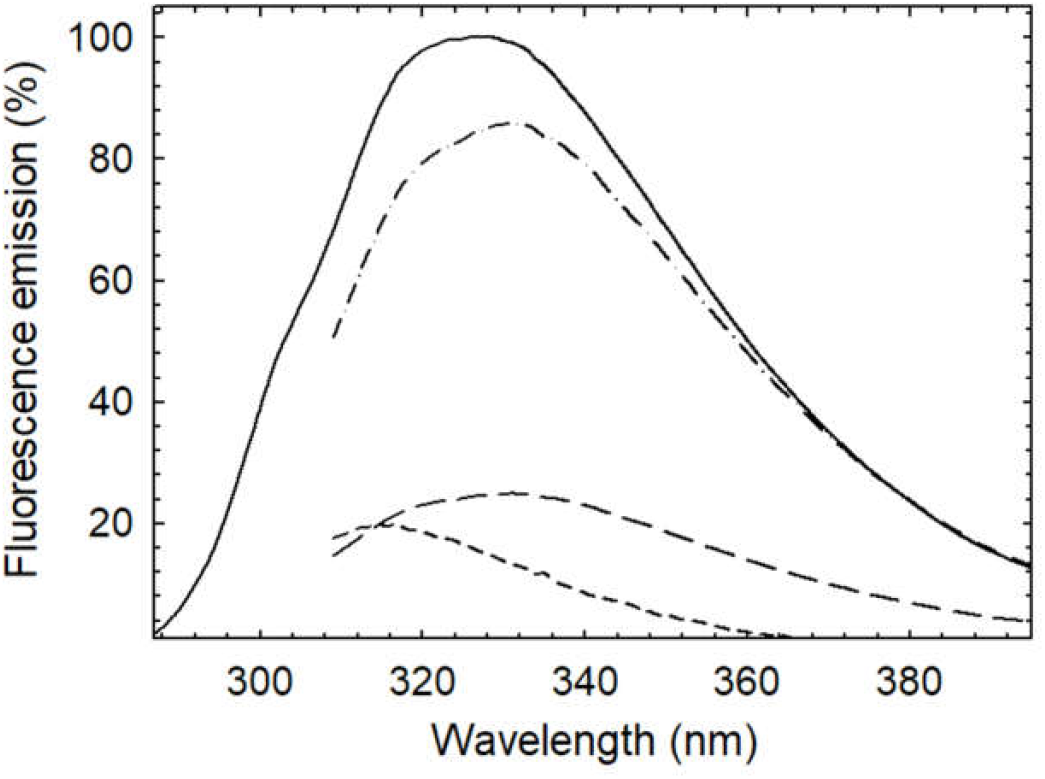
Fluorescence emission spectra of α-HL. Experimental spectra resulted from excitation at 275 nm (*solid line*) and 295 nm (*long dashed line*). This spectrum obtained upon excitation at 295 nm was normalized at wavelengths above 380 nm to obtain the Trp contributions (*dashed dot line*). Tyr contribution (*short-dashed line*) was calculated by subtracting the spectrum that represents the Trp contribution from that obtained upon excitation at 275 nm. Fluorescence emission units are arbitrary, expressed as percentages, and referred to the maximal value of α-HL upon excitation at 275 nm.

### 3.3. Hemolysis activity

The standard method used to measure the hemolytic activity of PFPs relies on recording the percentage of hemolysis observed after a reasonably long amount of time [37]. Thus, in order to assess the functional activity of purified α-HL, hemolytic assays were performed to analyze its pore-forming capacity at the plasmatic membrane (Figure 7). These assays employed erythrocytes from defibrinated horse and sheep blood, since these two animals show a marked species variation in the plasma membrane relative abundance of their respective phosphatidyl choline and sphingomyelin content [54], as it is shown in Table 2, while they contain a very similar amount of Chol in the order of 26% of total lipid species [54].

**Table 2:** Phospholipid distribution in erythrocytes from sheep and horse according to [54]. The substantial difference in PC content has been highlighted.

| *Phospholipid* | *Sheep* | *Horse* |
| --- | --- | --- |
| Phosphatidic acid | <0.3 | <0.3 |
| Phosphatidyl ethanolamine | 26.2 | 24.3 |
| Phosphatidyl serine | 14.1 | 18.0 |
| Phosphatidyl inositol | 2.9 | <0.3 |
| <u>Phosphatidyl choline</u> | <u>Undetectable</u> | <u>42.4</u> |
| Sphingomyelin | 51.0 | 13.5 |
| Lysophosphatidyl choline | Undetectable | 1.7 |
| Unidentified | 4.8 | Undetectable |

**Figure 7.**
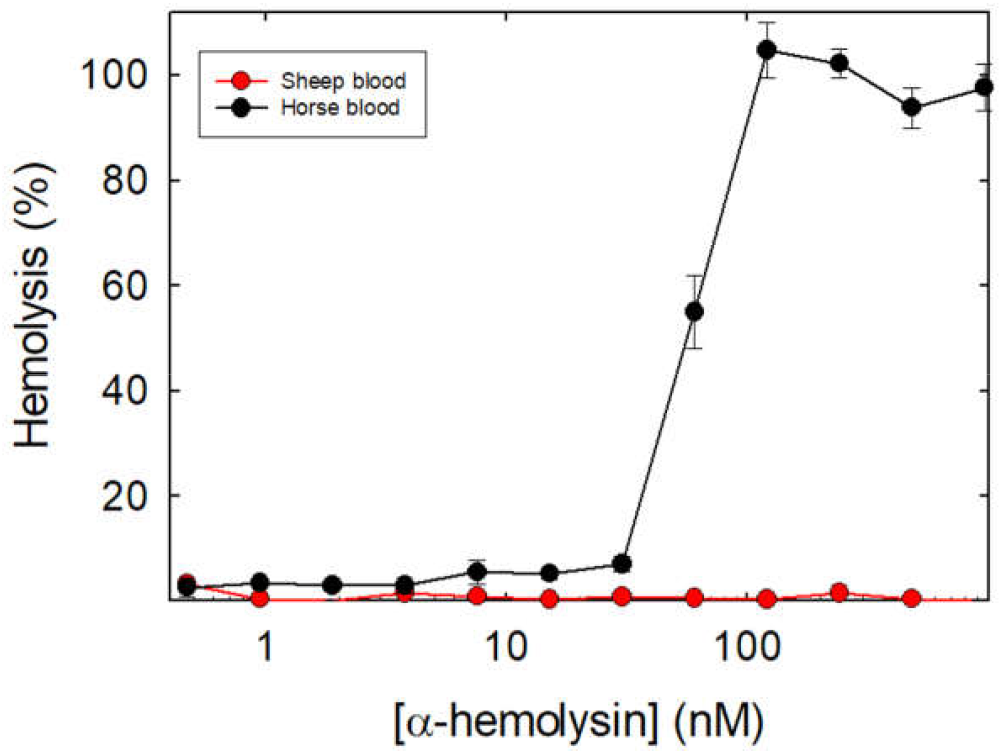
Hemolytic activity of the purified recombinant α-HL, expressed as a percentage of the hemolysis produced after 10 min of assay vs the logarithm of the protein concentration. The 100% value was calculated after the addition of Na_2_CO_3_ to a final concentration of 0.1 % (w/v).

The results obtained show that the protein was completely devoid of hemolytic activity when assayed against sheep blood even at protein concentrations where the same protein sample was able to produce 100 % of hemolysis against erythrocytes from horse blood (Figure 7). Overall, these results would emphasize the dramatic importance of the presence of phosphatidylcholine (PC) in the membrane for being recognized by α-HL. Indeed, they also showed that the purified recombinant protein was fully functional.

### 3.4. Binding to different lipid model vesicles

Erythrocytes, however, are complex systems that do not easily allow to draw conclusions about the nature of the lipid molecules involved in membrane binding and pore formation. As aforementioned, it seemed quite obvious from the hemolytic experiments that PC would play a key role in the activity of α-HL. However, the experiments performed with simpler systems, such as model lipid vesicles (large unilamellar vesicles; LUVs) of different lipidic compositions, revealed that this interpretation was simplistic.

ITC is a very suitable technique for studying the energetics of interactions between biological molecules [55], most especially when lipidic membranes are involved. Then, LUVs of four different compositions were prepared and used for these calorimetric experiments: DOPC, DOPC:SM (4:1), DOPC:Chol (4:1), and DOPC:SM:Chol (1:1:1). These lipid mixtures were chosen not only because they are systems thoroughly characterized in our laboratory [37, 39] but also because they roughly allow discrimination among the main components of horse erythrocytes, including its SM and high Chol content (in the order of 25 %) [54]. ITC results are shown in Figure 8 and Table 3.

**Table 3:** Thermodynamic parameters calculated for binding of purified recombinant α-HL to 100-nm-diameter LUVs of different lipidic composition studied by ITC.Results are the means of at least two independent experiments. Standard deviations are indicated. N is the number of lipid molecules involved in binding [56]. Kb is binding constant. Binding was not observed for DOPC and DOPC:SM (4:1) vesicles.

| | N | $K_b \cdot 10^4$<br>($M^{-1}$) | $\Delta G \cdot 10^2$<br>($kcal \cdot mol^{-1}$) | $\Delta H \cdot 10^4$<br>($kcal \cdot mol^{-1}$) | $\Delta S$<br>($cal \cdot mol^{-1} \cdot K^{-1}$) |
| --- | --- | --- | --- | --- | --- |
| **DOPC** | - | - | - | - | - |
| **DOPC:SM**<br>**4:1** | - | - | - | - | - |
| **DOPC:Chol**<br>**4:1** | $10,30 \pm 0,02$ | $9,45 \pm 4,29$ | $-71,6 \pm 2,90$ | $-1,59 \pm 0,59$ | $-41,50 \pm 0,93$ |
| **DOPC:SM:Chol**<br>**1:1:1** | $10,85 \pm 5,17$ | $16,45 \pm 8,72$ | $-8,93 \pm 0,41$ | $-1,86 \pm 1,24$ | $-19,75 \pm 8,77$ |

**Figure 8.**
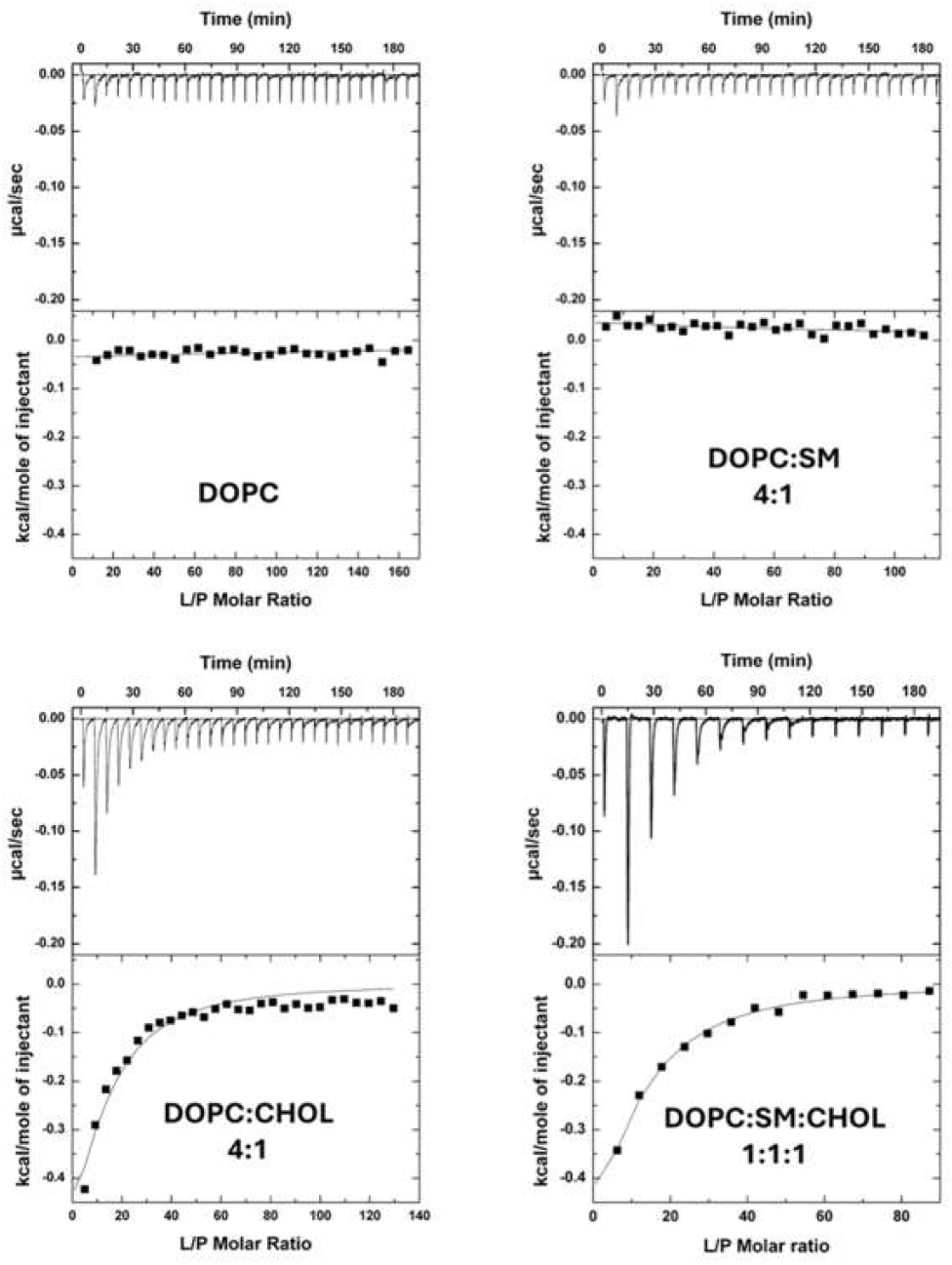
Binding of purified recombinant α-HL to 100-nm-diameter LUVs of different lipidic composition studied by ITC. Reactants’ concentrations were 5 µM αHL and 2.5 mM LUVs. Binding isotherms were adjusted to a model in which the protein binds the membrane involving N lipid molecules [56]. The c values (c=K×P_0_) were in the range 1–1000 (when binding was observed).

The most striking result was observing that, in the absence of any protein receptor, α-HL only bound to Chol-containing vesicles. Actually, the Chol content of these vesicles was very similar to that calculated for mammalian erythrocytes [54]. These results also agree with Chol acting as an enhancer of the interaction, as described before [20, 21]. On the other hand, in the conditions used for these experiments, the presence of SM seems irrelevant for the α-HL-membrane interaction, in agreement with the hemolysis experiments shown in Figure 7. It is also clear that the protein was unable to interact with plain homogeneous DOPC vesicles, enhancing not only the role of Chol but also of the presence of specific protein receptors at the plasma membrane for its cytolytic action.

### 3.5. SDS resistant oligomers in the presence of liposome vesicles

It is well documented that incubation of α-HL with liposomes of the adequate lipidic composition yields SDS-resistant oligomers, which seems to be directly related to the ability of this protein to make membrane pores [50]. Accordingly, the purified protein was able to assemble into SDS-resistant large oligomers, and this oligomerization was dependent on the incubation time, as shown in Figure 9.

**Figure 9.**
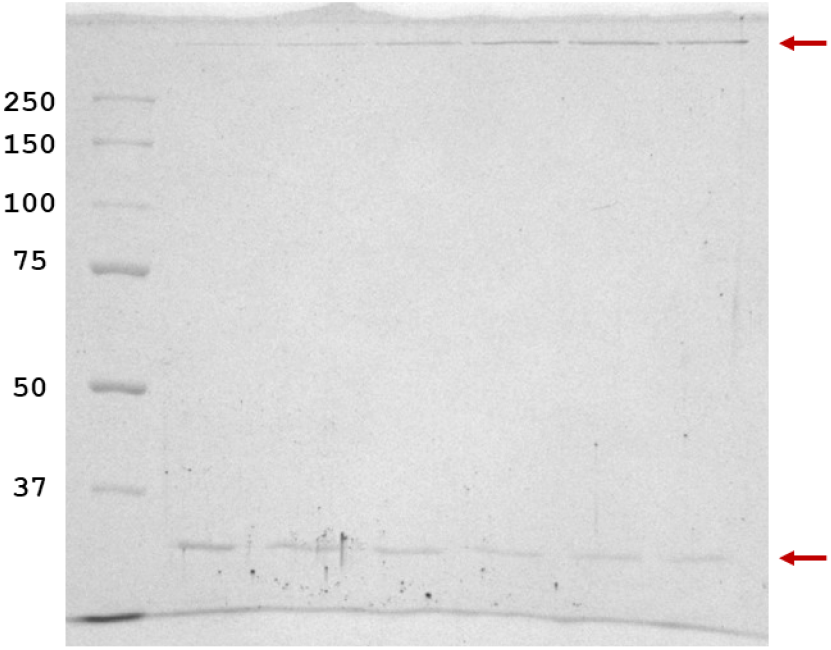
SDS-PAGE of a molecular weight protein standard (*left*) and the α-HL incubated with liposomes for various times (100, 300, 500, 700, 900 and 1100 seconds). The lower arrow marks the monomeric protein, while upper one corresponds to the SDS-resistant oligomers.

### 3.6. Reconstitution of a soluble transmembrane pore of α-HL on purified nanodisc water-soluble lipidic particles

In order to further explore the functionality of the purified α-HL, its ability to assemble into a pore was further analyzed. With this purpose, MSP1E3D1-DOPC:SM:Chol (1:1:1) purified empty nanodiscs were mixed with the protein at an α-HL/MSP1E3D1 molar ratio of 10/1. This assembly of individual pore water-soluble particles was made following this procedure to better mimic the real situation encountered by α-HL when integrating into the membrane of its target host. This mixture was then incubated at room temperature for 2 hours and analyzed by negative staining on a JOEL JMS 1400 electron microscope. The buffer employed was 20 mM Tris-HCl buffer, pH 7.4, 100 mM NaCl.

The negative staining electron microscopy image (Figure 10) shows multiple single particles corresponding to nanodiscs, most of which are compatible with the characteristic heptameric pore assembled structure of α-HL in membranes [9]. These results further prove there is no significant protein structural or functional changes affecting the recombinant purified α-HL herein reported and that nanodiscs represent a suitable water-soluble single-molecule platform in order to study α-HL functionality and to future in-built modifications aiming to develop different biotechnological tools.

**Figure 10.**
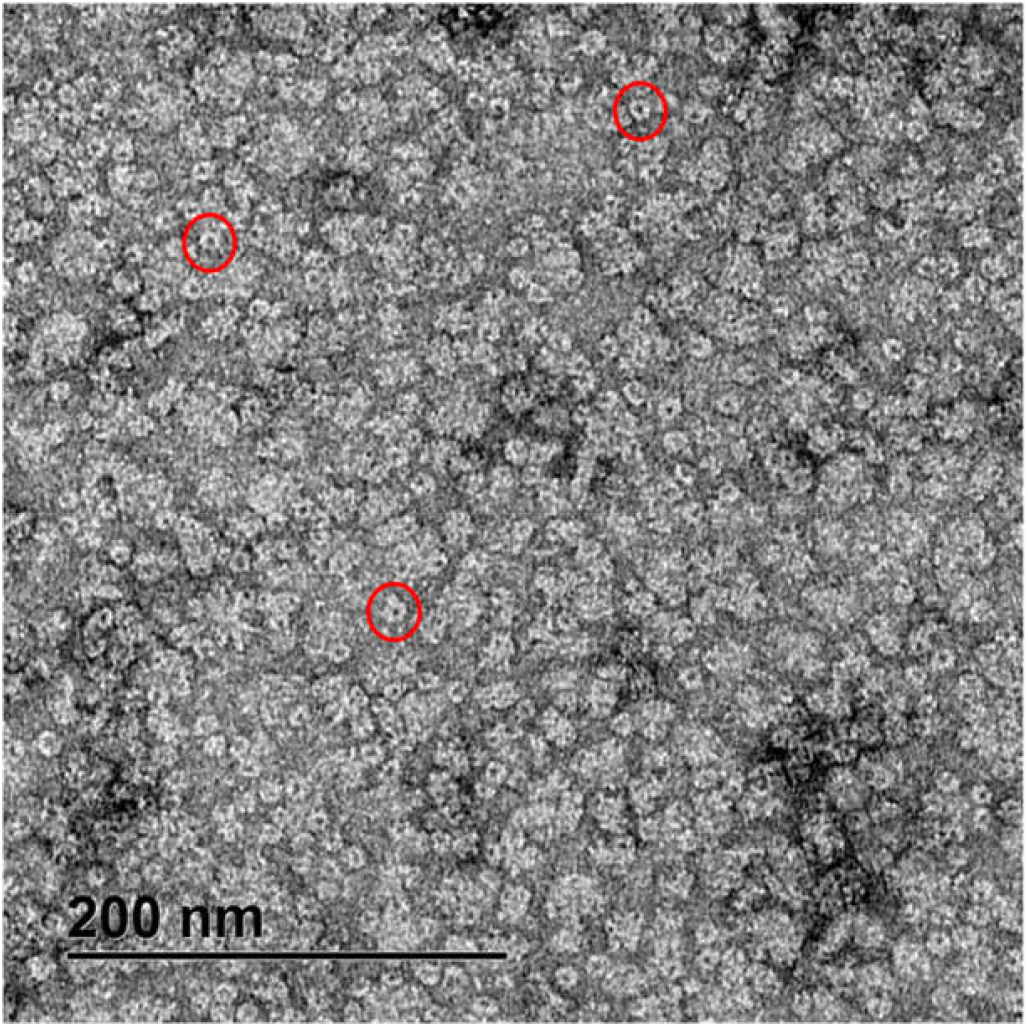
Negative staining electron microscopy micrograph of MSP1E3D1-DOPC:SM:Chol (1:1:1) purified nanodiscs after being incubated with α-HL. Three of these nanodiscs where pores are clearly observed have been circled (in red) as examples. Electron microscopy datasets were collected with JEOL JEM1400 transmission electron microscope and processed using DigitalMicrograph software (Gatan Inc.).

## 4. Conclusions

Overall, the results shown allow us to affirm that *S. aureus* α-HL has been produced and purified, following a simple protocol, in sufficient quantity to consider future biotechnological approaches. Functional and structural characterization has revealed that the protein is fully functional and confirms the key role played by Chol. It has also been shown that this purified toxin can be assembled in the form of individual pores within soluble lipid platforms such as those represented by Chol-containing nanodiscs.

## Abbreviations

CD: circular dichroism; Chol, cholesterol;
DOPC: dioleyl phosphatidylcholine;
HL: hemolysin;
ITC: isothermal titration calorimetry;
LUVs: large unilamelar vesicles;
PC: phosphatidylcholine;
PFP: pore-forming protein;
SM: sphingomyelin.

## 5. Acknowledgments

This work was financed by a REACT-EU grant from the Comunidad de Madrid to the ANTICIPA project of Complutense University of Madrid, UCM-Banco Santander Grants PR87/19-22556 and PR108/20-26896, and UnaEuropa (Unano) SF2106 (to A.M.-P.). Another UCM-Banco Santander Grant PR3/23-30816 (to S.G.-L.) was also part of this financing. D.H.-M. was supported by a *Ph D* appointment from Universidad Complutense de Madrid - Banco de Santander CT82/20-CT83/20. V.S. enjoyed an Erasmus traineeship grant from Università degli studi del Sannio, Benevento Italy (I BENEVEN 02).

